# AC-Diff: Anatomy–Contrast Disentangled Diffusion for Multi-Vendor Liver MRI Harmonization

**DOI:** 10.64898/2026.09.28.755033

**Authors:** Hansen Feng, Haotian Dang, Lei You, Hongyu Wang, Xiaobo Zhou

**Affiliations:** McWilliams School of Biomedical Informatics, The University of Texas Health Science Center at Houston; Tsinghua Shenzhen International Graduate School, Tsinghua University

## Abstract

Magnetic resonance imaging (MRI) exhibits substantial scanner and protocol dependent appearance variations, making harmonization challenging without distorting patient specific anatomy, particularly in abdominal imaging. We propose AC-Diff, an anatomy–contrast disentangled diffusion framework that formulates MRI harmonization as a factor-specific generative intervention. Using aligned multi-contrast supervision and cross-patient factor swapping, AC-Diff learns to separate anatomical structure from acquisition-dependent contrast. Unlike conventional image or latent diffusion, AC-Diff restricts stochastic generation to the contrast subspace while the source anatomy bypasses diffusion and is directly reused during reconstruction. Experiments on an in-house multi-vendor cohort and the external Duke Liver MRI dataset demonstrate improved target-domain alignment with strong structural preservation. In downstream liver segmentation, AC-Diff improves Dice from 0.942 for the original inputs to 0.954 for the harmonized images. These results support contrast-specific latent generation as a promising approach to anatomy-preserving MRI harmonization.

## 1 Introduction

Magnetic resonance imaging (MRI) provides rich soft-tissue contrast and is widely used for diagnosis, treatment planning, and quantitative image analysis. However, MRI appearance is highly sensitive to scanner vendor, field strength, pulse sequence, reconstruction pipeline, and acquisition protocol. These factors can substantially alter intensity distributions, tissue contrast, noise characteristics, and texture even when the underlying anatomy is unchanged. Consequently, models trained at one site may exploit scanner-specific signatures rather than biologically meaningful features, limiting their robustness across institutions.

This problem is particularly challenging in abdominal and liver MRI. The abdomen contains multiple adjacent organs with complex and deformable structures whose configurations vary with respiration, patient positioning, and acquisition conditions. At the same time, liver MRI exhibits substantial sequence and vendor dependent variation in tissue contrast and texture. Harmonization must therefore suppress acquisition-related appearance differences while preserving patient-specific morphology and internal structures. This creates a fundamental tension: stronger distribution alignment may improve cross-site consistency, while unconstrained generative translation can also modify anatomical content.

Existing harmonization methods address scanner variability through statistical correction, invariant representation learning, or image translation [Fortin et al., 2017, 2018, Dewey et al., 2019, Moyer et al., 2020, Dinsdale et al., 2021, Cackowski et al., 2023, Zhu et al., 2017, Choi et al., 2020, Liu and Yap, 2024, Beizaee et al., 2025]. Recent diffusion-based approaches further improve generative flexibility by conditioning harmonization on source anatomy and target-domain appearance [Wu et al., 2026, Scholz et al., 2025]. In particular, CACD [Scholz et al., 2025] learns disentangled anatomy and contrast representations that condition diffusion-based image generation. However, representation-level disentanglement or conditioning does not explicitly determine which factor is subjected to stochastic generation. This motivates a stronger architectural constraint on what the harmonization model is allowed to generate.

We therefore formulate harmonization as a factor-specific latent intervention. Stage A exploits geometrically aligned multi-contrast mDixon acquisitions, where Water, Fat, In-phase, and Opposed-phase images share the same underlying anatomy while exhibiting systematic contrast differences. These aligned observations provide structured supervision for learning anatomy and contrast factors. Cross-patient factor swapping further regularizes factor recombination by requiring source anatomy to remain compatible with contrast information obtained from a different patient.

Based on this formulation, we propose AC-Diff, an anatomy–contrast disentangled diffusion framework for multi-vendor liver MRI harmonization. Stage A partitions the spatial latent representation into anatomy and contrast components and regularizes them using multi-contrast supervision, contrastive learning, and cross-patient factor swapping. Stage B freezes the learned encoder and applies conditional diffusion exclusively to the contrast latent, while the source anatomy latent bypasses the stochastic process and is directly reused during decoding. This design preserves the source anatomy pathway outside the stochastic process and restricts diffusion to the contrast subspace.

We evaluate AC-Diff on an in-house multi-vendor liver MRI cohort and the external Duke Liver MRI dataset. Our evaluation examines target-domain alignment, preservation of source anatomy, and downstream liver segmentation. For the segmentation evaluation, we separately fine-tune the segmentation model from a common pretrained initialization on the original training images and on those produced by each harmonization method, using identical fine-tuning settings across all conditions. Together, these assessments examine whether contrast adaptation preserves anatomical information useful for downstream analysis.

The main contributions of this work are:

- We develop an anatomy–contrast representation learning framework using geometrically aligned multi-contrast mDixon acquisitions and cross-patient factor swapping to regularize separation of subject-specific anatomy and acquisition-dependent contrast.
- We formulate MRI harmonization as a factor-specific latent intervention and introduce contrast-only conditional diffusion, in which the source anatomy latent bypasses stochastic generation while only the contrast factor undergoes diffusion.
- We validate AC-Diff on an in-house multi-vendor liver MRI cohort and the external Duke Liver MRI dataset, reducing foreground Wasserstein distance by 46.2% and 19.3%, respectively, relative to the original inputs. Under standardized downstream fine-tuning, AC-Diff improves liver segmentation Dice from 0.942 to 0.954.

## 2 Related Work

### MRI harmonization

MRI harmonization aims to reduce scanner and protocol dependent variability while preserving hbiologically and clinically relevant information [Hu et al., 2023]. Statistical approaches such as ComBat have been widely used to remove site effects from derived imaging features, including diffusion and cortical-thickness measurements [Fortin et al., 2017, 2018]. At the image level, DeepHarmony learns a supervised mapping between scanner protocols using paired acquisitions [Dewey et al., 2019], while scanner-invariant representation learning and adversarial unlearning seek to suppress acquisition-specific information in learned features [Moyer et al., 2020, Dinsdale et al., 2021]. Unpaired image translation further relaxes the requirement for cross-scanner correspondences; CycleGAN [Zhu et al., 2017] and multi-domain translation models such as StarGAN v2 [Choi et al., 2020] have therefore been widely adopted as generic harmonization baselines. More recent approaches include VAE–GAN-based self-supervised harmonization such as ImUnity [Cackowski et al., 2023] and source-free distribution alignment using normalizing flows [Beizaee et al., 2025]. Despite their flexibility, image-level translation objectives do not directly specify which anatomical or acquisition-related factors are permitted to change during harmonization.

### Anatomy–contrast disentanglement

Disentangled representations provide a more explicit mechanism for separating anatomical content from acquisition-dependent appearance. CALAMITI uses information-bottleneck principles to learn distinct anatomy and contrast representations for unsupervised MR harmonization [Zuo et al., 2021], while HACA3 extends this idea with contrast, anatomy, and artifact aware attention for multi-site harmonization across heterogeneous MR acquisitions [Zuo et al., 2023]. Related approaches use multi-contrast supervision, randomized contrast transformations, or domain invariant objectives to encourage separation between structural and acquisition factors [Dewey et al., 2020, Cackowski et al., 2023, Moyer et al., 2020]. These methods demonstrate the value of explicitly modeling anatomy and appearance; however, representation level factorization alone does not determine which latent variables are actually modified by the subsequent generative process.

### Diffusion-based harmonization

Diffusion models provide stable high-fidelity generation through iterative denoising [Ho et al., 2020, Song et al., 2021], and latent diffusion reduces computational cost by moving generation into a learned representation space [Rombach et al., 2022]. Durrer et al. [2024] adapt DDPMs to paired MRI contrast harmonization while explicitly incorporating source structural information. More recently, HCLD performs unpaired volumetric MRI harmonization using conditional latent diffusion with source anatomy and target style constraints [Wu et al., 2026]. The work most closely related to ours is CACD [Scholz et al., 2025], which combines a diffusion autoencoder with supervised contrastive learning and domain-agnostic contrast augmentation to obtain anatomy and contrast conditioning representations. These representations jointly condition DDIM-based image generation. In contrast, AC-Diff learns spatial anatomy–contrast factors from geometrically aligned real multi-contrast acquisitions and cross-patient factor swapping, and restricts the stochastic generation process itself to the contrast latent. The source anatomy latent bypasses diffusion and is directly reused during decoding. Thus, the key distinction of AC-Diff is not only how anatomy and contrast are represented, but which factor is allowed to undergo stochastic generation.

## 3 Method

### 3.1 Problem Formulation and Framework Overview

Given an unpaired source image *x*^i,s^ and a target-domain reference *x*^j,t^, we define the harmonized image as

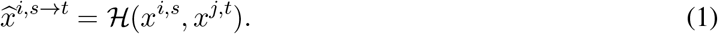

The objective is to align the distribution of harmonized images with the target domain while preserving the patient-specific anatomy of *x*^i,s^. No spatial or pixel-wise correspondence between the two inputs is assumed.

Because acquisition-dependent contrast and anatomy are entangled in MRI, direct image translation may inadvertently modify anatomical structures. We therefore formulate harmonization as a factor-specific latent intervention. Stage A learns an anatomy–contrast factorization from geometrically aligned multi-contrast mDixon acquisitions. Stage B freezes the learned encoder and performs conditional diffusion only in the contrast subspace, while the source anatomy representation bypasses the generative process. The preserved anatomy and generated target-domain contrast are recomposed and decoded into the harmonized image.

### 3.2 Learning an Anatomy–Contrast Representation

Let *x*^i,A^ denote an image of patient *i* under mDixon contrast *A ∈{*W, F, IP, OP*}* . These four contrasts are geometrically aligned within each acquisition, providing structured supervision for learning anatomy and contrast-dependent representations.

A four-level U-Net VAE encoder produces *z ∈* R^16*×*16*×*16^, which is partitioned channel-wise as

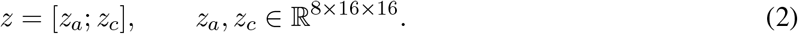

Here, *z*_a_ and *z*_c_ represent anatomy and acquisition-dependent contrast, respectively. Each factor is passed through a projection head consisting of global average pooling followed by a linear mapping to obtain 128-dimensional embeddings, *e*_a_ = *P*_a_(*z*_a_) and *e*_c_ = *P*_c_(*z*_c_). The contrast embedding *e*_c_ is additionally used for FiLM modulation in the decoder.

#### Factor-level contrastive supervision

We impose complementary invariances using the aligned multi-contrast data. For an anchor (*i, A*), the positive sets are

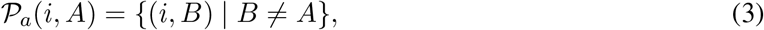

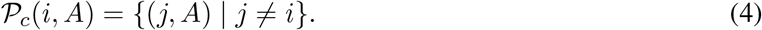

Thus, anatomy embeddings are matched across contrasts of the same patient, whereas contrast embeddings are matched across patients acquired with the same contrast. We use a temperature-scaled multi-positive contrastive objective [Chen et al., 2020, Khosla et al., 2020]:

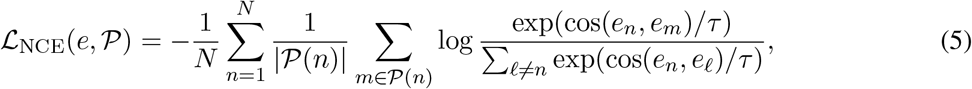

where cos(*·, ·*) denotes cosine similarity and *τ* is the temperature. The factorization objective is

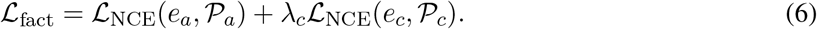

#### Cross-patient factor swapping

Embedding-level invariance does not preclude the spatial contrast latent *z*_c_ from retaining patient-specific anatomical information. A within-patient swap would provide limited pressure against such leakage, because anatomical information carried by the contrast latent would remain consistent with the reconstruction target. We therefore deliberately pair anatomy and contrast representations from different patients.

For *i ≠ j*, the anatomy latent from *x*^i,A^ is combined with the contrast latent from *x*^j,B^:

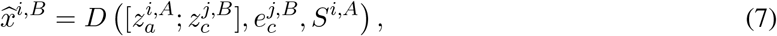

where *S*^i,A^ denotes source-derived skip features. The reconstruction target is the geometrically aligned image *x*^i,B^ of patient *i*. Hence, donor-specific anatomy from patient *j* is inconsistent with the supervision target, discouraging reliance on anatomical information in the contrast pathway without assuming perfect disentanglement.

The target contrast embedding modulates decoder features through FiLM:

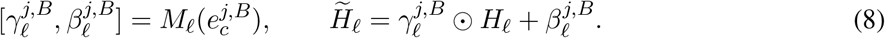

**Stage-A objective**. For the cross-patient reconstruction in Eq. (7), we use

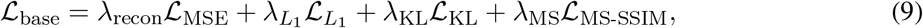

where the image-space terms are evaluated between 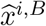 and *x*^i,B^, and 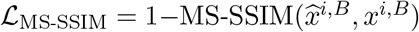. The complete Stage-A objective is

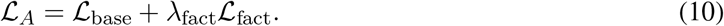

Together, the contrastive and cross-patient reconstruction objectives regularize the anatomy–contrast factorization used by Stage B.

### 3.3 Contrast-Space Conditional Diffusion for Harmonization

As shown in Figure 1, Stage B implements the factor-specific intervention by applying conditional diffusion exclusively to the contrast latent. For unpaired source and target-domain images *x*^i,s^ and *x*^j,t^, the frozen Stage-A encoder provides the source anatomy latent 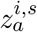 and the target contrast latent 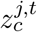 with corresponding embeddings 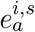 and 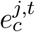 .

**Figure 1:**
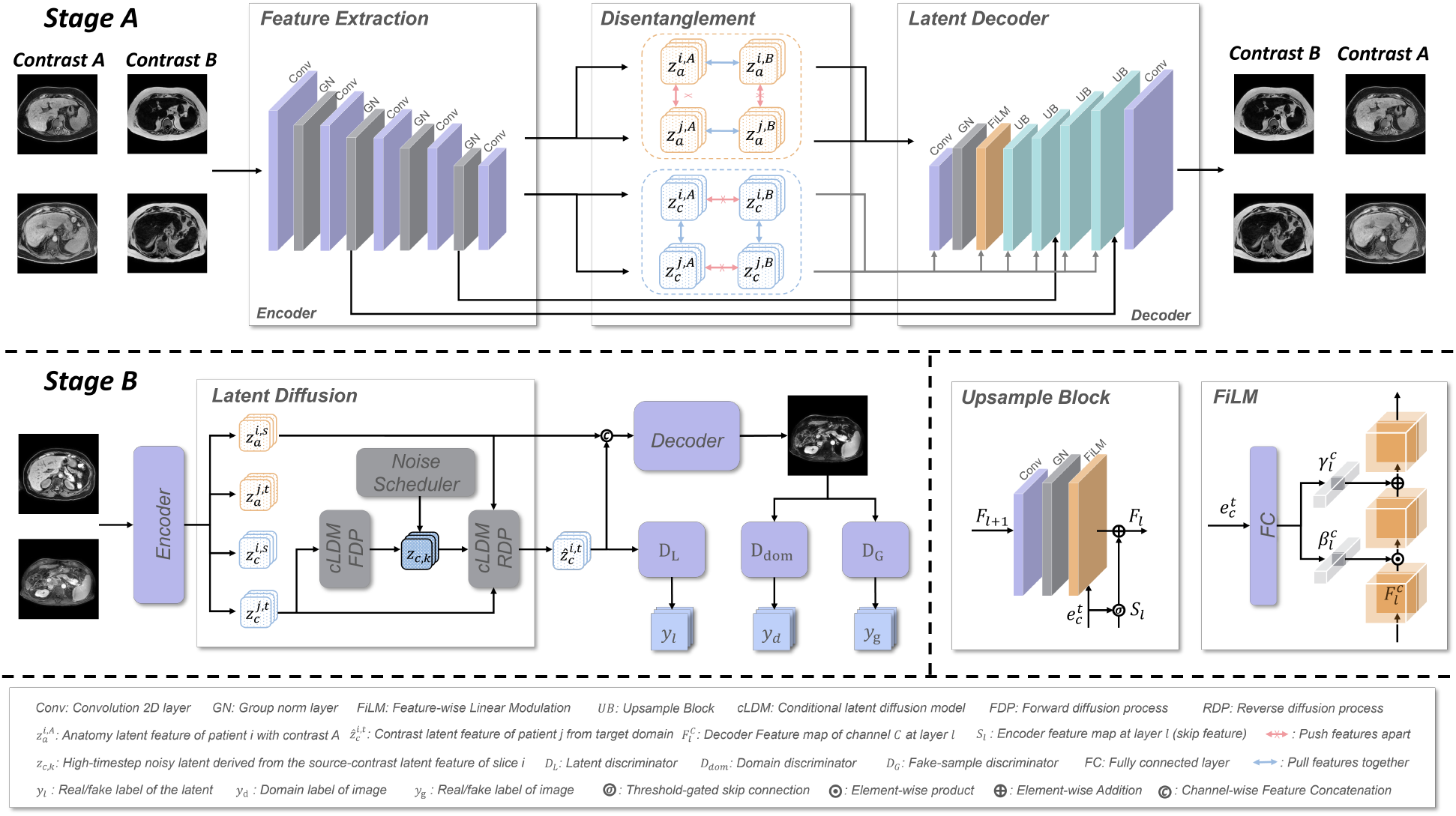
Overview of AC-Diff. Stage A learns anatomy and contrast representations from aligned multicontrast MRI using contrastive supervision and cross-patient factor swapping. Stage B freezes the encoder and applies conditional diffusion only to the target contrast latent, while the source anatomy latent bypasses the stochastic process and is reused during decoding.

The source anatomy and target contrast embeddings are mapped to conditioning tokens

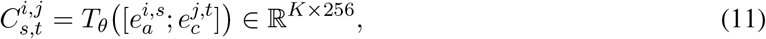

which condition the latent U-Net through multi-scale cross-attention. For brevity, we denote 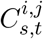 by *C* below. During training, the clean diffusion target is 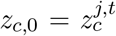 and Gaussian noise is applied only to this contrast latent:

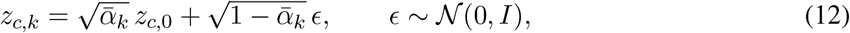

where 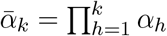 The denoiser is optimized using

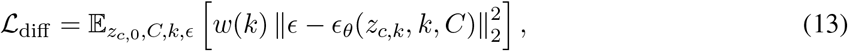

where *w*(*k*) applies Min-SNR-*γ* weighting [Hang et al., 2023] with additional emphasis on low-noise states; its exact form is given in Appendix A.1. Since *x*^i,s^ and *x*^j,t^ are sampled independently, no paired cross-domain image supervision is required.

For latent- and image-space regularization, we recover a differentiable estimate of the clean contrast latent,

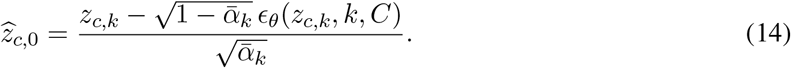

The predicted contrast latent is then recombined with the source anatomy and decoded:

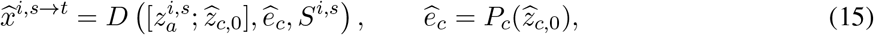

where *S*^i,s^ denotes source-derived skip features. Thus, 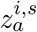 bypasses the stochastic process and is directly reused during reconstruction, while diffusion operates only on the contrast component.

At inference, 50-step DDIM sampling generates the contrast latent from Gaussian noise. For consistent harmonization across subjects, the individual target embedding 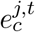is replaced by the population-level target representation *e*^⋆^ defined in Appendix A.2.

### 3.4 Stage-B Training Objective

The contrast-space diffusion objective in Section 3.3 learns to generate a target-domain contrast latent conditioned on the source anatomy and target contrast context. However, matching the latent denoising objective alone does not ensure that the decoded image exhibits the desired target-domain appearance, nor that the intended anatomy and contrast factors remain consistent after decoding. We therefore augment *L*_diff_ with three complementary constraints: distribution alignment, factor consistency, and training stabilization. The Stage-B generator is optimized with

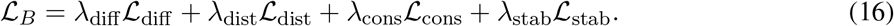

#### Distribution alignment

The diffusion loss acts in contrast-latent space, whereas harmonization is ultimately evaluated in the decoded image space. We therefore align the generated sample with the target domain at both levels:

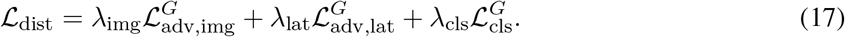

Here, 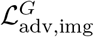 encourages the decoded harmonized image 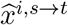 to follow the target-domain image distribution, while 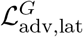 aligns the predicted clean contrast latent 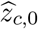 with real target contrast latents. An auxiliary acquisition classification loss 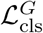 further encourages the generated image to exhibit the requested target acquisition label *y*_t_.

The corresponding discriminators are trained with standard binary adversarial objectives. The image discriminator uses the raw target domain MRI *x*^j,t^ as real and 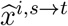 as fake, while the latent discriminator an acquisition-classification head supervised by the source and target domain labels. uses 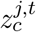 and 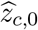as real and fake contrast latents, respectively. T he image discriminator additionally includes

#### Factor consistency

Distribution alignment alone does not guarantee that the decoded image retains the intended factor composition. We therefore re-encode 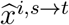 using the frozen Stage-A encoder and recover its anatomy and contrast embeddings, 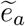 and 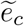 The consistency objective is

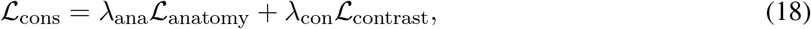

where *L*_anatomy_ encourages 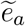 to remain consistent with the source anatomy 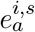, and *L* _contrast_ encourages 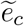 to match the target contrast representation 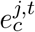 This provides a closed-loop constraint on the intended source-anatomy/target-contrast recomposition after decoding.

#### Training stabilization

Finally, we regularize cases in which the source image already belongs to the target domain. In this setting, harmonization should introduce minimal appearance change. We therefore use

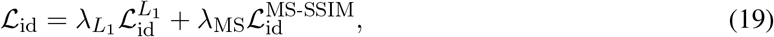

to preserve image intensity and structure. When the pretrained Stage-A decoder is fine-tuned during the later stage of training, we additionally apply a reconstruction anchor *L*_dec_ to limit decoder drift:

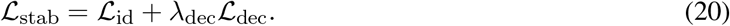

Together, these objectives extend contrast-space diffusion from latent generation to image-level harmonization: distribution alignment promotes target-domain appearance, factor consistency preserves the intended source-anatomy with target-contrast composition, and stabilization limits unnecessary changes during decoder adaptation.

## 4 Experiment

### 4.1 Datasets

#### In-house multi-vendor cohort

Our in-house cohort contains 109 MRI series from 18 unique patients acquired across GE, Philips, and Siemens scanners. The data are split into 80 training, 19 validation, and 10 test series. Geometrically aligned multi-contrast mDixon acquisitions, including Water (W), Fat (F), In-phase (IP), and Opposed-phase (OP) images, are used in Stage A to provide structured supervision for anatomy–contrast factorization. Detailed acquisition statistics are provided in Appendix A.3.

#### Duke Liver MRI dataset

We use the Duke Liver MRI Dataset [Macdonald et al., 2023] for external evaluation and downstream liver segmentation. For downstream liver segmentation, we select category B (axial precontrast fat-suppressed T1-weighted MRI), comprising 85 cases from 84 patients. We adopt a fixed patient-disjoint split of 42 training, 13 validation, and 29 test patients.

Duke data are excluded from AC-Diff training, while segmentation annotations are used exclusively for downstream fine-tuning and evaluation. Further details are provided in Appendix A.4.

### 4.2 Harmonization Evaluation

#### Image-space evaluation

We compare AC-Diff with ComBat [Fortin et al., 2018], CycleGAN [Zhu et al., 2017], StarGAN v2 [Choi et al., 2020], Harmonizing Flows [Beizaee et al., 2025], and DDPM [Ho et al., 2020]. Table 1 highlights that high source similarity does not necessarily imply target-domain alignment. DDPM achieves the highest MS-SSIM among harmonization methods, yet its Wasserstein distance on Duke exceeds that of the original images (0.191 vs. 0.181). In contrast, AC-Diff achieves the lowest Wasserstein distance on our cohort and Duke (0.113 and 0.146, respectively), together with the highest PSNR and gradient similarity among harmonization methods on both datasets. These results indicate a favorable balance between source preservation and target-domain alignment, including on the external Duke dataset, which is not used to train the harmonization model.

**Table 1:** Quantitative comparison of MRI harmonization methods on our multi-vendor liver MRI cohort and the Duke Liver MRI dataset. Results are reported as patient-level means. Harm. Flows denotes Harmonizing Flows. SSIM denotes multi-scale SSIM (MS-SSIM). MS-SSIM, PSNR, and gradient similarity evaluate preservation of source image structure and local detail (↑), whereas the foreground Wasserstein distance measures intensity-distribution discrepancy from the target real Philips mDixon Water images (*↓*). Best results are shown in bold.

| Method | Our Liver Dataset |  |  |  | Duke Liver Dataset |  |  |  |
| --- | --- | --- | --- | --- | --- | --- | --- | --- |
| | SSIM ↑ | PSNR ↑ | Grad. ↑ | Wass. ↓ | SSIM ↑ | PSNR ↑ | Grad. ↑ | Wass. ↓ |
| Original | 1.000 | ∞ | 1.000 | 0.210 | 1.000 | ∞ | 1.000 | 0.181 |
| ComBat | 0.760 | 19.998 | 0.967 | 0.222 | 0.848 | 18.437 | 0.965 | 0.168 |
| CycleGAN | 0.695 | 20.718 | 0.956 | 0.165 | 0.736 | 22.358 | 0.965 | 0.165 |
| StarGAN v2 | 0.527 | 14.188 | 0.939 | 0.160 | 0.827 | 19.976 | 0.969 | 0.152 |
| Harm. Flows | 0.760 | 21.460 | 0.970 | 0.272 | 0.922 | 21.609 | 0.971 | 0.156 |
| DDPM | **0.903** | 23.055 | 0.975 | 0.145 | **0.964** | 26.149 | 0.978 | 0.191 |
| AC-Diff (ours) | 0.891 | **25.054** | **0.979** | **0.113** | 0.940 | **26.762** | **0.981** | **0.146** |

#### Factor-space analysis

To further examine this balance, we evaluate all methods using the frozen Stage-A encoder (Figure 2). AC-Diff combines low target-centroid contrast distance with the highest source–output anatomy similarity, whereas several baselines show negative changes in target-centroid distance. These observations are consistent with the intended anatomy–contrast separation, although this model-dependent analysis provides supporting evidence rather than an independent measure of anatomical fidelity.

**Figure 2:**
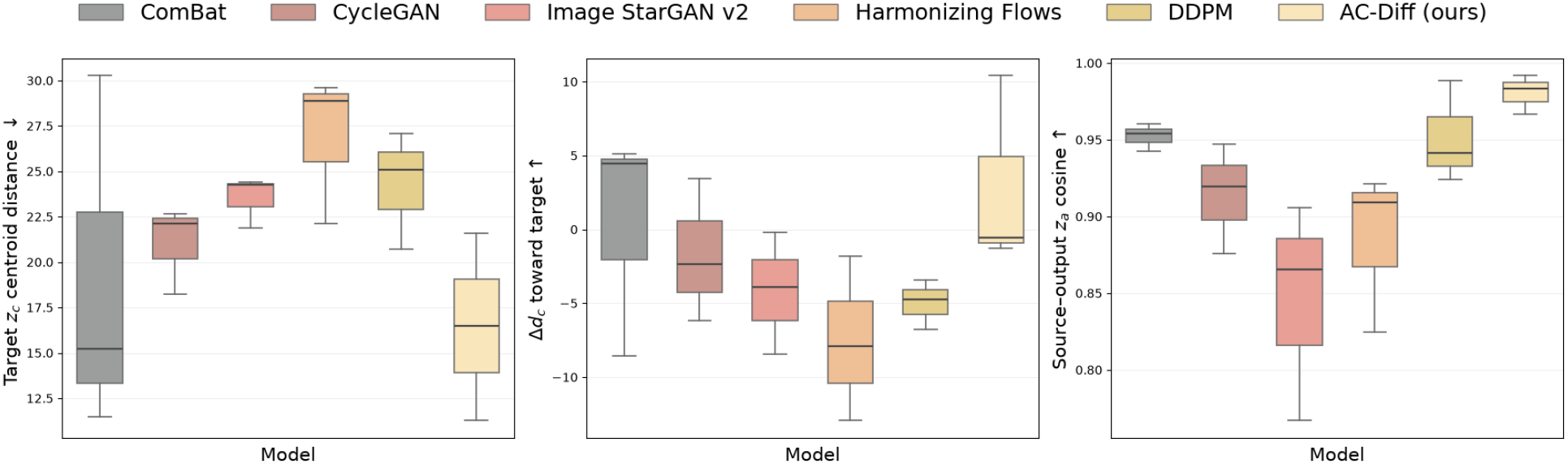
Patient-level factor-space analysis on our multi-vendor liver MRI cohort using the frozen Stage-A encoder as a common evaluator. Each box summarizes patient-level mean values over non-target-domain cases, with the center line denoting the median. **Left:** distance between the output contrast representation *z*_c_ and the centroid of held-out Philips/mDixon-W target representations (↓). **Middle:** change in target-centroid distance, 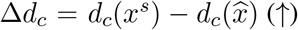, where positive values indicate movement toward the target contrast distribution. **Right:** cosine similarity between source and output anatomy representations *z*_a_ (*↑*), measuring preservation of anatomy-related information.

### 4.3 Downstream Liver Segmentation

To evaluate the downstream utility of harmonized images, we independently fine-tune the same pretrained TotalSegmentator MRI model [Akinci D’Antonoli et al., 2025] for each harmonization method. We use the Duke category-B subset with a fixed patient-disjoint split of 42 training, 13 validation, and 29 test patients. All methods, including the original non-harmonized baseline, share the same initialization and fine-tuning configuration. Segmentation annotations are used exclusively for downstream training and evaluation, rather than harmonization training.

As shown in Table 2, AC-Diff achieves the highest observed mean Dice (0.954) and recall (0.960), compared with 0.942 and 0.943 for the original images, respectively. Relative to DDPM, AC-Diff improves Dice from 0.943 to 0.954 while reducing HD95 from 6.518 to 4.856 mm and ASSD from 1.553 to 1.361 mm. The original images exhibit an elevated mean HD95, driven by a small number of cases with substantial boundary errors, despite their relatively high Dice. Qualitative results in Figure 3 further illustrate the segmentation predictions obtained after method-specific fine-tuning, highlighting the differences in liver delineation across harmonization methods.

**Table 2:** Downstream liver segmentation on the Duke category-B subset after method-specific fine-tuning of TotalSegmentator MRI. Results are patient-level means over 29 held-out test patients. Abs. denotes absolute relative volume error; HD95 and ASSD are reported in millimeters. Best results are shown in bold.

| Input | Dice ↑ | Precision ↑ | Recall ↑ | Abs. ↓ | HD95 (mm) ↓ | ASSD (mm) ↓ |
| --- | --- | --- | --- | --- | --- | --- |
| Original | 0.942 | 0.941 | 0.943 | 0.024 | 20.570 | 3.916 |
| ComBat | 0.942 | 0.946 | 0.940 | 0.027 | 6.973 | 1.776 |
| CycleGAN | 0.873 | 0.866 | 0.896 | 0.132 | 28.288 | 5.983 |
| StarGAN v2 | 0.935 | 0.939 | 0.932 | 0.030 | 7.541 | 1.836 |
| Harmonizing Flows | 0.940 | 0.946 | 0.934 | 0.027 | 9.379 | 2.027 |
| DDPM | 0.943 | 0.947 | 0.939 | 0.022 | 6.518 | 1.553 |
| AC-Diff (ours) | **0.954** | **0.949** | **0.960** | **0.015** | **4.856** | **1.361** |

**Figure 3:**
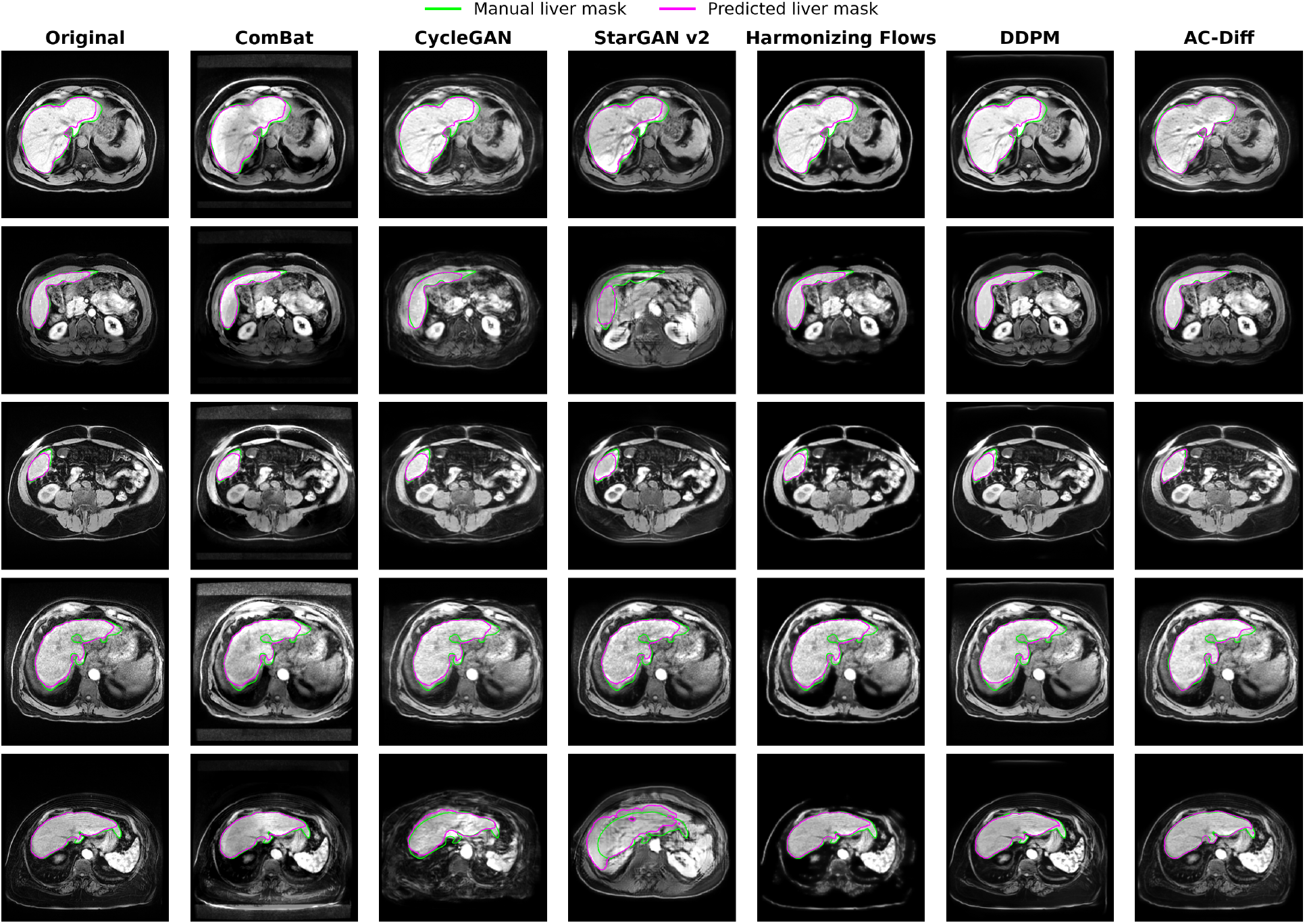
Qualitative comparison of downstream liver segmentation on the Duke Liver MRI dataset. Columns show original images and images harmonized by different methods. Green contours indicate manual liver annotations, and magenta contours indicate predictions from TotalSegmentator MRI models separately fine-tuned for each image condition from a common pretrained initialization using identical fine-tuning settings.

### 4.4 Ablation Study

Table 3 shows that diffusion scope affects the balance between source preservation and target-domain alignment. The no-diffusion variant directly combines the source anatomy latent with a target-domain contrast latent through factor swapping and decodes the resulting representation without diffusion-based generation. With Stage-A factorization and a frozen decoder, full-latent diffusion improves source similarity over this direct-swap baseline but worsens target alignment. Restricting diffusion to *z*_c_ improves both under matched settings, increasing MS-SSIM from 0.737 to 0.796 and reducing Wasserstein distance from 0.142 to 0.121. Decoder fine-tuning further improves all three metrics, whereas removing adversarial regularization increases Wasserstein distance from 0.113 to 0.164 despite retaining relatively high source similarity.

**Table 3:** Ablation study of AC-Diff on our multi-vendor liver MRI cohort, examining Stage-A factorization, diffusion scope, adversarial regularization (Adv.), and decoder fine-tuning. MS-SSIM and PSNR are computed relative to the source images; Wasserstein distance is computed against real Philips mDixon Water images. Best results are shown in bold.

| Stage A | $z_c$ Diff. | Full Diff. | Adv. | Unfreeze Decoder | MS-SSIM ↑ | PSNR (dB) ↑ | Wass. ↓ |
| --- | --- | --- | --- | --- | --- | --- | --- |
| × | × | ✓ | ✓ | ✓ | 0.419 | 9.873 | 0.263 |
| ✓ | × | × | ✓ | × | 0.647 | 17.825 | 0.129 |
| ✓ | × | ✓ | ✓ | × | 0.737 | 19.079 | 0.142 |
| ✓ | ✓ | × | ✓ | × | 0.796 | 20.492 | 0.121 |
| ✓ | ✓ | × | × | ✓ | 0.866 | 24.381 | 0.164 |
| ✓ | ✓ | × | ✓ | ✓ | **0.891** | **25.054** | **0.113** |

## Discussion

Our results suggest that restricting harmonization to acquisition-related contrast provides a better trade-off between target-domain alignment and anatomical preservation than unconstrained image translation. Although DDPM achieves slightly higher source–output MS-SSIM, AC-Diff obtains stronger target alignment, higher gradient similarity, and improved downstream liver segmentation. This supports the central design of AC-Diff: rather than regenerating the full image representation, diffusion is confined to the contrast subspace while source anatomy bypasses the stochastic process.

Several limitations remain. Anatomy–contrast disentanglement is not perfect, and residual anatomical information in the contrast representation may lead to local deformation during diffusion. Harmonization is also sensitive to foreground occupancy, since variations in field of view, cropping, and body size can affect intensity distributions and learned contrast representations. In addition, the current model operates slice-wise and relies on aligned multi-contrast mDixon data for Stage-A supervision. Extending AC-Diff to volumetric harmonization, stronger anatomy-consistency constraints, and foreground-aware modeling represents an important direction for future work.

## 6 Conclusion

We presented AC-Diff, an anatomy–contrast disentangled diffusion framework for multi-vendor liver MRI harmonization. By restricting diffusion to the contrast representation and reusing the source anatomy latent during decoding, AC-Diff adapts image appearance while limiting direct generative changes to anatomical features. Experiments on an in-house multi-vendor cohort and the external Duke Liver MRI dataset demonstrate improved target-domain alignment with strong structural preservation, supported by representation-space analysis and improved downstream liver segmentation under a common fine-tuning protocol. These findings support contrast-specific generation as a practical approach to balancing appearance adaptation and anatomical fidelity.

## Acknowledgments

This project is supported by the CPRIT-funded project (GRANT ID RP250043).

## A Experiment Settings

### A.1 Implementation Details

#### Data preprocessing

DICOM pixel values were first converted using the rescale slope and intercept. Within each MRI series, unreadable slices and inconsistent scout/localizer-like images were excluded, and the remaining slices were ordered by spatial position. Intensities were normalized independently for each 3D series: values were clipped to the 1st and 99th percentiles of the series-wide intensity distribution and linearly mapped to [*−* 1, 1].

To preserve the native aspect ratio, each *H × W* slice was symmetrically padded along its shorter dimension with a background value of *−*1, forming a square image that was isotropically resized to 256 *×*256. During inference, the harmonized output was resized to the padded native dimensions and cropped to recover the original *H × W* image geometry.

#### Architecture

Stage A operates on 256 *×* 256 single-channel slices. The VAE latent comprises 16 channels at 16 *×* 16 spatial resolution, equally partitioned into eight anatomy and eight contrast channels. Both factors are projected into 128-dimensional global embeddings. Gated additive skip connections are enabled at decoder levels 2 and 4.

In Stage B, the concatenated anatomy–contrast condition is mapped to *K* = 8 tokens of width 256. Diffusion uses *T* = 1000 noise levels with a linear variance schedule from *β*_1_ = 10^*−*4^ to *β*_T_ = 2 × 10^*−*2^. Inference employs 50-step DDIM sampling.

#### Diffusion weighting

We use Min-SNR-*γ* weighting [Hang et al., 2023] with an additional emphasis on low-noise diffusion states. For timestep *k*, the signal-to-noise ratio is

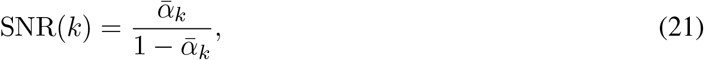

and the Min-SNR weight for noise prediction is

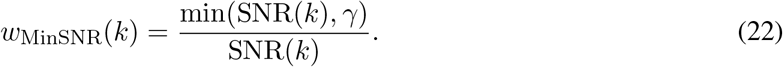

We additionally apply a low-noise boost

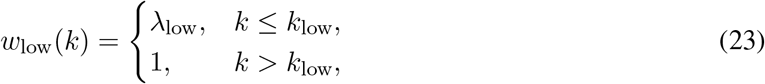

giving the final diffusion weight

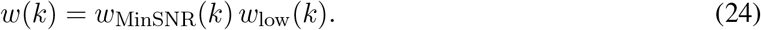

We use *γ* = 5.0, *k*_low_ = 20, and *λ*_low_ = 2.0 in all main experiments.

#### Training schedule

The Stage-A encoder remains frozen throughout Stage B. The latent adversarial objective is activated after 2000 optimization steps and applied only to low-noise states (*k ≤* 20). We ob-erved that activating this objective from the beginning could destabilize optimization, occasionally causing gradient explosion before the diffusion model had learned a meaningful target contrast-latent distribution. Delaying the adversarial objective and restricting it to low-noise states mitigated this instability.

After epoch 20, the decoder is fine-tuned with a learning rate two orders of magnitude lower than that of the diffusion generator. A reconstruction anchor is retained to stabilize decoder updates.

#### Downstream segmentation fine-tuning

We fine-tune a pretrained TotalSegmentator MRI model independently for each harmonization method using the Duke category-B subset (axial precontrast fat-suppressed T1-weighted MRI). A fixed patient-disjoint split assigns 42, 13, and 29 patients to training, validation, and testing, corresponding to 43, 13, and 29 cases, respectively. All methods, including the original non-harmonized baseline, use the same pretrained initialization, patient partition, 20 training epochs, and initial learning rate of 10^*−*3^. Each fine-tuned model is evaluated on the corresponding images from the held-out test patients. Segmentation labels are used exclusively for downstream fine-tuning and evaluation, not for training the harmonization models.

### A.2 Robust Target Representation and Geometry-Preserving Inference

A single reference image may encode patient- or slice-specific appearance that is not representative of the target acquisition domain. We therefore construct a population-level target contrast representation. Contrast embeddings are first averaged across central slices within each series and then across series from the same patient, yielding patient-level representations 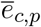 These are aggregated using the geometric median:

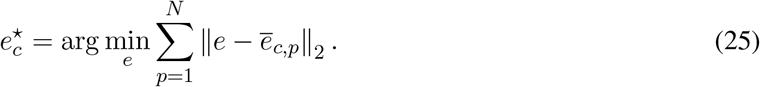

The geometric median is estimated using Weiszfeld iterations, giving each patient equal influence regardless of the number of available slices or series. The resulting 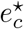 is used as a fixed target-domain representation for all source images during population-reference inference.

To preserve native image geometry, rectangular slices are symmetrically padded to a square field of view before isotropic resizing to the network input resolution. After harmonization, the output is resized back to the padded native extent and cropped to the original *H × W* dimensions. This avoids aspect-ratio distortion while retaining the original spatial geometry.

### A.3 In-House Dataset Details

The cohort is highly imbalanced across acquisition groups. Philips/mDixon-W constitutes the dominant domain, whereas Philips/THRIVE and Siemens/Dixon-W contain only a single patient and Siemens/VIBE contains relatively few patients. This imbalance motivated the use of Philips/mDixon-W as the target domain, for which a substantially larger number of subjects and series is available. It also limits reliable patient-disjoint evaluation for sparsely represented acquisition groups.

Patient counts in Table 4 are reported independently for each acquisition group and are therefore not additive, since the same patient may contribute multiple protocols or series. The cohort contains 18 unique patients after deduplication. Data splitting is performed at the patient level to avoid subject overlap between training, validation, and test sets. For acquisition groups represented by only a single patient, a patient-disjoint validation or test subset cannot be formed; such groups are therefore used only where permitted by the training split and are not reported as independent held-out domains.

**Table 4:** Composition of the in-house multi-vendor MRI cohort.

| Acquisition group | Patients | Series | Slices |
| --- | --- | --- | --- |
| GE/LAVA | 8 | 18 | 1,932 |
| Philips/THRIVE | 1 | 12 | 1,836 |
| Philips/mDixon-W | 10 | 61 | 7,177 |
| Siemens/Dixon-W | 1 | 1 | 88 |
| Siemens/VIBE | 4 | 17 | 1,384 |
| Total | 18<sup>†</sup> | 109 | 12,417 |

The complete collection contains 141 MRI series. We retain 109 series belonging to the five predefined harmonization groups and exclude the remaining 32 series from the main harmonization analysis according to protocol-based selection criteria.

### A.2 Duke Liver Dataset Details

We use category B of the Duke Liver MRI Dataset [Macdonald et al., 2023], comprising axial precontrast fat-suppressed T1-weighted MRI with liver segmentation annotations. The selected subset contains 85 annotated series from 84 patients. we partition the subset into training, validation, and testing sets at the patient level (Table 5). All series from the same patient are assigned to a single partition to prevent patient overlap across subsets.

**Table 5:**
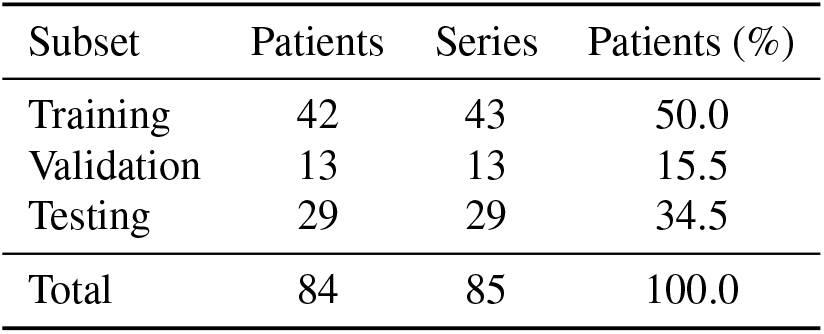
Patient-disjoint partition of the Duke category-B subset for downstream liver segmentation. Percentages denote the proportion of patients in each subset.

We select this category for its relevance to the Philips mDixon Water target domain and its suitability for downstream evaluation. Both fat-suppressed T1-weighted images and mDixon water-only images reduce the contribution of fat signal, providing a more comparable contrast setting than a mixture of fat-suppressed and non-fat-suppressed acquisitions. Restricting the evaluation to precontrast images also avoids additional heterogeneity associated with contrast-enhancement phases. This selection provides a controlled setting for assessing harmonization and its impact on liver segmentation, without assuming that conventional fat suppression and Dixon water–fat separation produce identical image contrast.

### A.5 Evaluation Metrics

#### Harmonization metrics

We evaluate harmonization using multi-scale structural similarity (MS-SSIM), peak signal-to-noise ratio (PSNR), gradient similarity, and foreground Wasserstein distance. MS-SSIM, PSNR, and gradient similarity are computed between the source and harmonized images to assess structural and local-detail preservation. Foreground Wasserstein distance measures the discrepancy between the foreground intensity distributions of harmonized images and real Philips mDixon Water images, which serve as the target-domain reference. Higher MS-SSIM, PSNR, and gradient similarity, and lower Wasserstein distance indicate better performance under their respective evaluation criteria.

#### Downstream segmentation metrics

We evaluate liver segmentation using Dice, precision, recall, absolute relative volume error (Abs.), 95th percentile Hausdorff distance (HD95), and average symmetric surface distance (ASSD).

Let *P* and *G* denote the predicted and ground-truth foreground voxel sets, respectively. The absolute relative volume error is

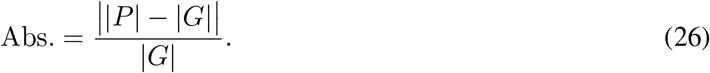

To evaluate boundary accuracy, let *∂P* and *∂G* denote the surfaces of the predicted and ground-truth segmentations. The shortest Euclidean distance from a point *x* to a surface *A* is *d*(*x, A*) = min_a*∈*A_∥ *x − a*∥ _2_. The 95th percentile Hausdorff distance is

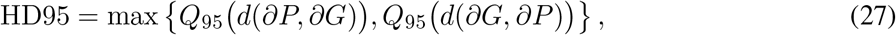

where *d*(*∂P, ∂G*) denotes the set of point-to-surface distances from *∂P* to *∂G*, and *Q*_95_ denotes the 95th percentile.

The average symmetric surface distance is

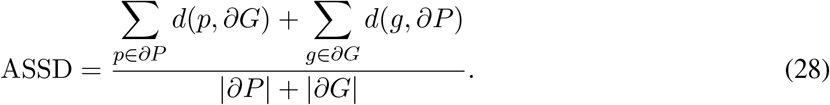

Surface distances are computed in physical space and reported in millimeters. Lower Abs., HD95, and ASSD indicate smaller volume and boundary errors.

### B Stage-A Qualitative Factor-Swapping Results

Figure 4 qualitatively illustrates the factor-recombination behavior learned in Stage A. Each row shows geometrically aligned mDixon contrasts from the same source patient, including Water (W), Fat (F), Inphase (IP), and Opposed-phase (OP) images. For the translated results, the source anatomy latent is retained while the contrast latent is replaced by a W-contrast latent sampled from a different patient.

**Figure 4:**
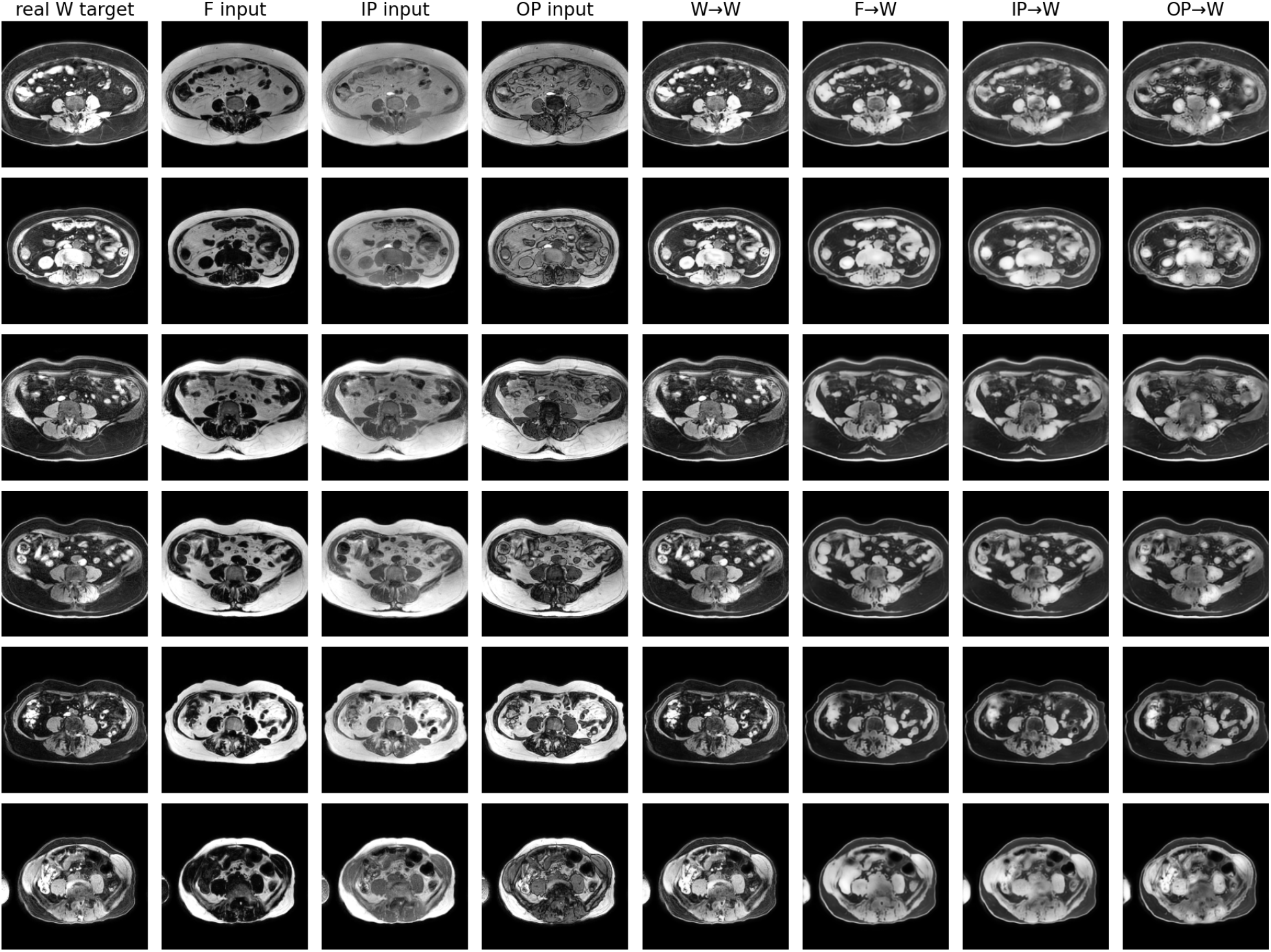
Qualitative Stage-A cross-patient contrast-latent swapping. Each row corresponds to one source patient with geometrically aligned mDixon Water (W), Fat (F), In-phase (IP), and Opposed-phase (OP) images. The leftmost column shows the real W image of the source patient. For W→W, F→W,IP→W, and OP→W, the anatomy representation is taken from the corresponding source image, while theW-contrast latent is provided by a different patient. The translated images exhibit W-like appearance while largely retaining the source patient’s anatomical configuration.

Across the examples, F*→* W, IP*→* W, and OP*→* W outputs consistently shift toward the appearance of the real W image despite substantial differences among the input contrasts. At the same time, the overall body contour and major abdominal structures remain spatially consistent with the source subject. The W *→*W results provide a same-domain reference and show only limited changes when the requested contrast already matches the source domain. These observations provide qualitative evidence that the learned representation supports cross-patient recombination of anatomy and contrast. Importantly, they demonstrate functional factor separation at the decoded-image level rather than proving that the spatial contrast latent is entirely free of anatomical information.

### C Stage-B Qualitative Results

Figure 5 presents qualitative comparisons of Stage-B harmonization results on representative slices from our multi-vendor liver MRI cohort. We compare AC-Diff with ComBat, CycleGAN, StarGAN v2, Harmonizing Flows, and DDPM.

**Figure 5:**
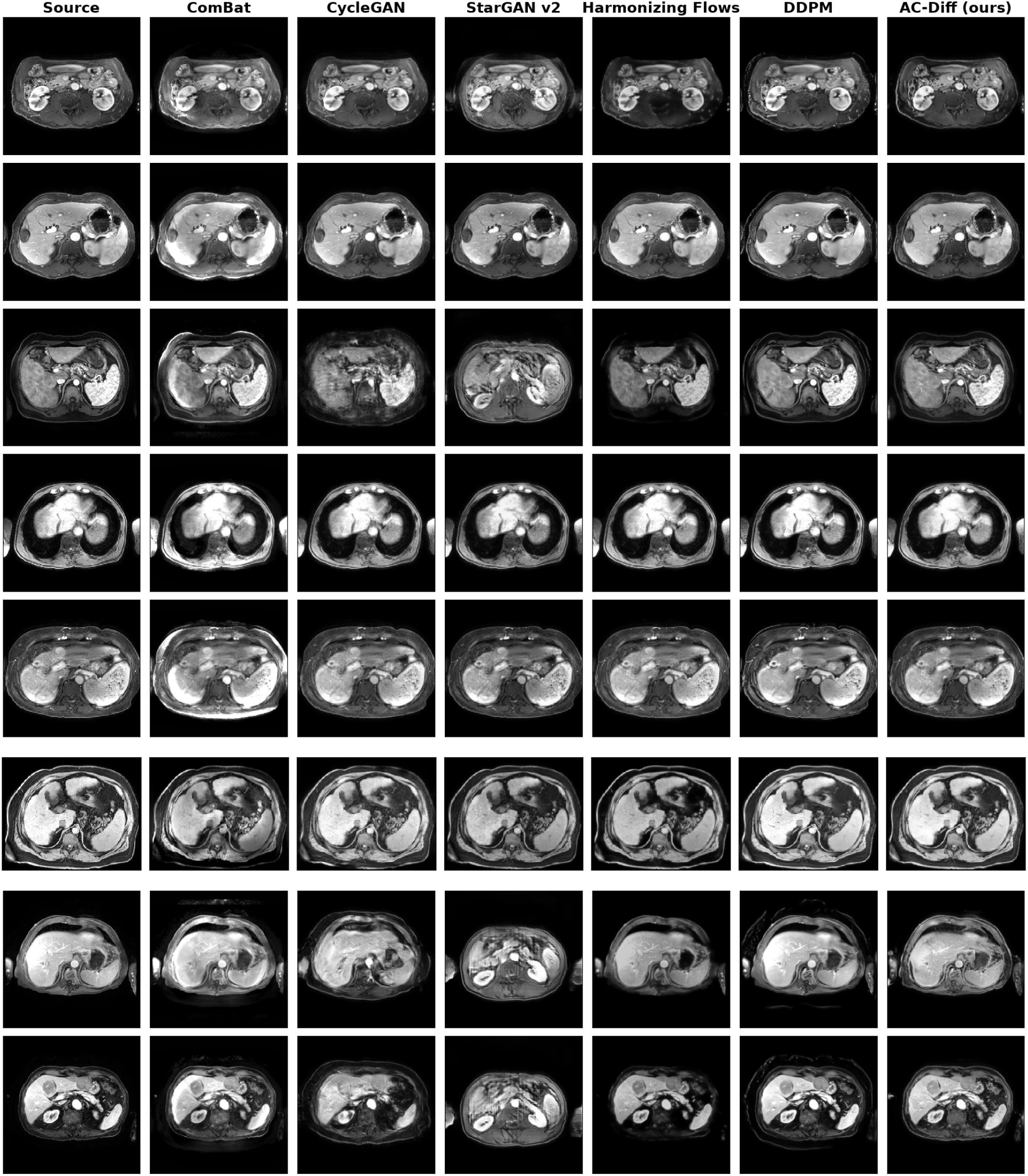
Qualitative comparison of Stage-B harmonization results. Each row shows one representative source slice from our multi-vendor liver MRI cohort and the corresponding outputs produced by different harmonization methods. AC-Diff achieves visually consistent target-style harmonization while largely preserving the source anatomical configuration. In contrast, ComBat mainly changes global intensity statistics, CycleGAN may introduce blurring or local appearance distortion, StarGAN v2 exhibits severe structural artifacts, and Harmonizing Flows and DDPM, although more stable, remain less consistent than AC-Diff in balancing appearance adaptation and anatomical preservation.

**Figure 6:**
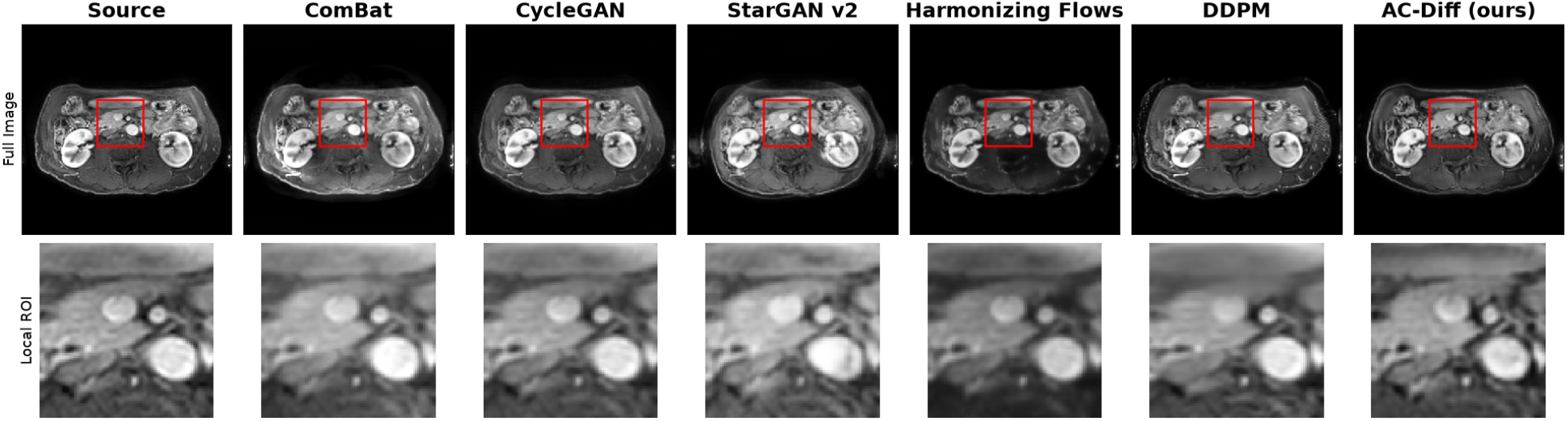
Local structural comparison of Stage-B harmonization methods. The top row shows the full source image and harmonized outputs, with the same anatomical region highlighted in red. The bottom row presents magnified views of the corresponding ROI. The zoomed comparison facilitates inspection of local boundaries, vessel morphology, and fine structural consistency after harmonization. StarGAN v2 exhibits pronounced local distortion, whereas AC-Diff retains local structures that remain visually consistent with the source anatomy while adapting image appearance.

Overall, AC-Diff produces the most balanced results between target-domain appearance adaptation and anatomical preservation. Across the examples, AC-Diff consistently adjusts image contrast and intensity style while maintaining the overall abdominal contour, organ layout, and internal structural boundaries of the source image. ComBat mainly performs global intensity normalization and preserves gross anatomy, but its harmonization effect is comparatively limited. CycleGAN occasionally moves toward the target appearance, but in some cases introduces local blurring or loss of fine structural detail. StarGAN v2 shows clear instability, with severe structural distortion and unrealistic texture patterns in some particular examples. Harmonizing Flows and DDPM generally yield more plausible outputs than the adversarial baselines, but still exhibit residual appearance shifts or reduced structural fidelity in challenging cases.

These qualitative observations are consistent with the quantitative results in the main text: AC-Diff provides strong target-domain alignment while avoiding the pronounced anatomical degradation observed in less constrained image-translation baselines.

### D Baseline Training Difficulties and Evaluation Limitations

We additionally attempted to apply CACD and HCLD, two methods developed for brain MRI harmonization, to our abdominal MRI task. Neither attempt yielded a successful harmonization model under the configurations we evaluated. We describe these unsuccessful attempts below to explain their absence from the quantitative comparison. These observations concern our implementations and training settings and should not be interpreted as evidence that either method is inherently unsuitable for abdominal MRI.

#### D.1 CACD Training Difficulties

We adapted the public CACD implementation to our abdominal MRI pipeline through a dataset adapter, using the same fixed patient split and preprocessing as the other baselines. We trained a 2D configuration based on its identity-preserving DiffAE variant with single-channel 256 *×* 256 slices and scanner labels specifying the target domain. To investigate training sensitivity, we tested different combinations of GIN-based random convolution perturbations, random bias-field corruption, gamma contrast transformations, geometric augmentation, and random resizing. We also varied the batch size over *{*2, 4, 6, 8*}*to alter the composition of positive and negative examples during training. Across the tested configurations, qualitative inspection revealed persistent anatomical degradation: source-to-target outputs were dominated by noise-like patterns, while reconstructions were overly smooth or intensity-distorted and failed to faithfully retain abdominal structures.

As these attempts did not yield a reliable harmonization model, we did not include CACD in the quantitative comparison. The observed difficulties may relate to the greater variability in organ configuration, tissue appearance, and field of view in abdominal imaging, which could complicate anatomy–appearance disentanglement. However, our experiments do not isolate the primary cause or rule out optimization and implementation issues. We therefore report this outcome as an unsuccessful adaptation under the tested configurations, rather than evidence of an inherent limitation of CACD, and acknowledge the resulting gap in baseline coverage.

In contrast, AC-Diff uses geometrically aligned multi-contrast mDixon acquisitions to supervise disentanglement, providing anatomically corresponding views with different contrasts that may help preserve anatomical information during representation learning.

#### D.2 HCLD Stage-B Output Degeneration

We implemented HCLD as a two-stage volumetric pipeline, comprising 3D autoencoder training followed by conditional latent diffusion training using the learned autoencoder. Without pretrained abdominal MRI weights, we trained the autoencoder and diffusion model from scratch on our in-house abdominal MRI training set. Stage A was trained for 30 epochs and produced anatomically recognizable reconstructions. Stage B was configured for 80 epochs with a batch size of 1 and gradient accumulation over four iterations.

The Stage-B objective combined diffusion denoising, latent content preservation, Gram-matrix style matching, adversarial style alignment, and gradient consistency:

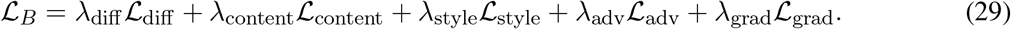

The diffusion and gradient-consistency terms used L2 losses, and the adversarial style term was activated after a five-epoch burn-in. The initial weights in our implementation were (*λ*_diff_, *λ*_content_, *λ*_style_, *λ*_adv_, *λ*_grad_) = (1, 10, 10, 1, 10000). After inspecting the loss magnitudes in TensorBoard, we tested a manually reweighted configuration, (1, 5, 1000, 2, 500). Both configurations used the same burn-in and DDIM settings: 50 training steps, 30 forward validation steps, and 10 reverse validation steps. This reweighting did not resolve the structural degeneration observed in the monitored Stage-B outputs.

For one monitored case at epoch 29, the logged source, target, and reconstruction mean intensities were 0.1513, 0.1408, and 0.0247, respectively. Although these scalar statistics alone do not establish anatomical failure, the corresponding validation images showed extensive loss of recognizable abdominal structure. Both the reconstruction and source-to-target synthesis outputs degenerated into localized high-intensity regions, indicating a failure beyond a simple contrast mismatch.

A possible contributing factor is the difficulty of balancing style alignment and anatomical preservation when adapting a brain-oriented framework to heterogeneous abdominal anatomy. Style-related objectives may permit substantial spatial distortion when content constraints are insufficient, even after loss reweighting. We encountered similar instability during the development of AC-Diff, suggesting that this behavior is not unique to HCLD. However, our experiments do not isolate the contribution of individual loss terms or fully rule out implementation and sampling issues. We therefore did not include HCLD in the quantitative comparison and report this outcome as an unsuccessful training attempt under our evaluated configurations. The absence of a reliable HCLD baseline remains a limitation of our comparative evaluation.

### E Theoretical Analysis of Contrast-Subspace Diffusion

#### E.1 Conditional Diffusion Objective in the Contrast Latent Space

Stage B models the conditional distribution of target-domain contrast latents while keeping the source anatomy latent outside the diffusion process. Let *z*_c,0_ denote a clean target contrast latent and *c* the conditioning information derived from the source anatomy and target contrast context. With *α*_k_ = 1 *− β*_k_ and 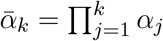 the forward process is

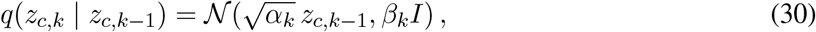

which gives

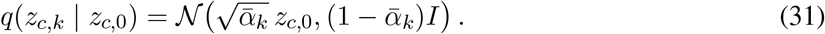

Equivalently,

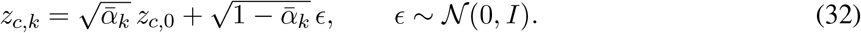

For the variational formulation, consider a Gaussian reverse process

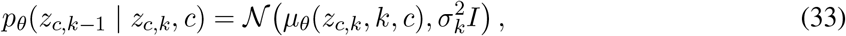

with fixed positive variances and prior *p*(*z*_c,T_) = *N* (0, *I*). For each (*z*_c,0_, *c*), the conditional negative log-likelihood satisfies

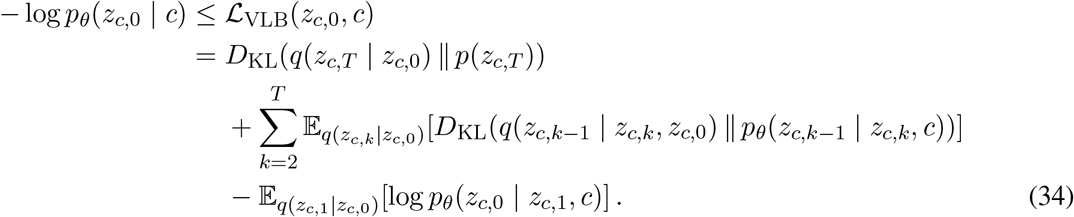

The expectations are over the forward process conditional on the clean latent; the bound does not hold pointwise for an arbitrary sampled trajectory.

Under the standard DDPM noise-prediction parameterization, the intermediate Gaussian KL terms reduce, up to parameter-independent constants, to timestep-weighted noise-prediction errors. In AC-Diff, we optimize the surrogate objective

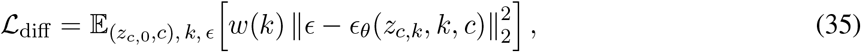

where (*z*_c,0_, *c*) follows the training sampling procedure, *ϵ ∼ N* (0, *I*), and *k* is sampled from the training timestep distribution. The weight *w*(*k*) is the Min-SNR weight with the low-noise boost specified in Appendix A.1. This practical weighting differs from the coefficients of the exact variational bound; consequently, *L*_diff_ is not itself claimed to upper-bound the conditional negative log-likelihood. The additional Stage-B regularizers are also separate from this variational formulation.

The diffusion objective specifies how the contrast-latent distribution is learned, but does not ensure anatomical invariance. Stage A uses aligned multi-contrast supervision, factor-level contrastive learning, and cross-patient factor swapping to encourage functional separation of anatomy and contrast. These constraints do not guarantee that the spatial contrast latent is free of anatomical information. The global projection *P*_c_(*z*_c_) provides contrast conditioning but does not remove the direct spatial *z*_c_ pathway to the decoder. Thus, Stage B restricts generation to the learned contrast coordinates, whose anatomical independence remains an empirical property.

#### E.2 Error Bound for Contrast-Subspace Diffusion

We analyze the propagation of latent errors through a fixed decoder. For a given source image, hold its skip features *S*^s^ fixed and define

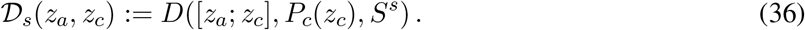

This definition includes both the spatial contrast pathway and its FiLM-mediated effects. To isolate diffusion scope, the comparison below uses the same decoder parameters and source skip features for both models. Let

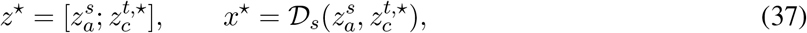

where 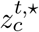 is a desired target-domain contrast latent. Here, *x*^⋆^ is a decoder-space reference rather than an observed paired ground-truth image; any mismatch between this reference and an ideal harmonized image is outside the latent-error bounds below.

Consider full-latent and contrast-subspace outputs

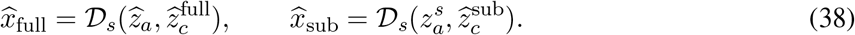

Assume that *D*_s_ is block-wise Lipschitz over the relevant latent region:

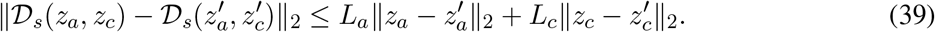

Then

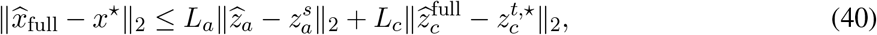

whereas

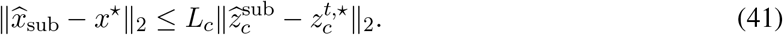

Reusing 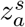 therefore eliminates the explicit anatomy-latent error term from this bound. However, the two models can have different contrast errors, and comparing upper bounds alone does not establish an ordering of their actual output errors.

For a local analysis, assume differentiability at *z*^⋆^ and sufficiently small latent perturbations. Let *J*_a_ and *J*_c_ denote the corresponding Jacobians of *D*_s_, and define

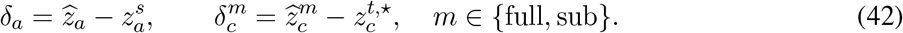

To first order,

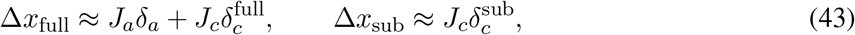

where 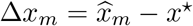 For the fixed source and reference, take expectations over generation randomness and assume finite second moments. Write 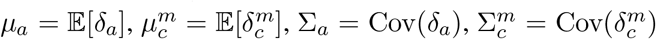 and 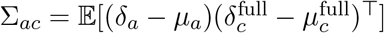. The resulting second-moment expressions are

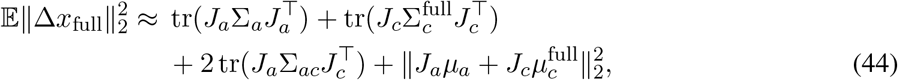

and

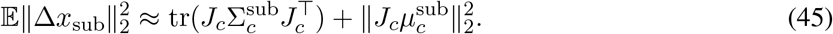

Under the additional assumptions of zero-mean errors, S_ac_ = 0, and matched contrast-error covariances 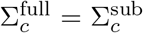, the full-latent expression contains the additional nonnegative term tr 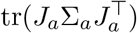 . This conditional comparison explains the benefit of avoiding direct anatomy-latent perturbations; it does not establish these assumptions for the trained models or guarantee lower image-space error in general.

Finally, image-space error is not an anatomy-specific metric. Residual anatomical information in *z*_c_ can still affect decoded structures through *J*_c_, even when 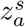 is unchanged. Decoder fine-tuning also changes the mapping and its sensitivities, which the fixed-decoder comparison does not cover. The analysis therefore motivates contrast-subspace generation as an architectural constraint, while the quality of factor separation and anatomical preservation must be assessed empirically.

## Notes

### Competing Interest Statement

The authors have declared no competing interest.

## References

Jean-Philippe Fortin, Drew Parker, Birkan Tunç, Takanori Watanabe, Mark A. Elliott, Kosha Ruparel, David R. Roalf, Theodore D. Satterthwaite, Ruben C. Gur, Raquel E. Gur, Robert T. Schultz, Ragini Verma, and Russell T. Shinohara. Harmonization of multi-site diffusion tensor imaging data. NeuroImage, 161:149–170, 2017. doi: 10.1016/j.neuroimage.2017.08.047.

Jean-Philippe Fortin, Nicholas Cullen, Yvette I. Sheline, Warren D. Taylor, Irem Aselcioglu, Philip A. Cook, Phil Adams, Crystal Cooper, Maurizio Fava, Patrick J. McGrath, Melvin McInnis, Mary L. Phillips, Madhukar H. Trivedi, Myrna M. Weissman, and Russell T. Shinohara. Harmonization of cortical thickness measurements across scanners and sites. NeuroImage, 167:104–120, 2018. doi: 10.1016/j.neuroimage.2017.11.024.

Blake E. Dewey, Can Zhao, Jacob C. Reinhold, Aaron Carass, Kathryn C. Fitzgerald, Elias S. Sotirchos, Shiv Saidha, Jiwon Oh, Dzung L. Pham, Peter A. Calabresi, Peter C. M. van Zijl, and Jerry L. Prince. Deepharmony: A deep learning approach to contrast harmonization across scanner changes. Magnetic Resonance Imaging, 64:160–170, 2019. doi: 10.1016/j.mri.2019.05.041.

Daniel Moyer, Greg Ver Steeg, Chantal M. W. Tax, and Paul M. Thompson. Scanner invariant representations for diffusion mri harmonization. Magnetic Resonance in Medicine, 84(4):2174–2189, 2020. doi: 10.1002/mrm.28243.

Nicola K. Dinsdale, Mark Jenkinson, and Ana I. L. Namburete. Deep learning-based unlearning of dataset bias for mri harmonisation and confound removal. NeuroImage, 228:117689, 2021. doi: 10.1016/j.neuroimage.2020.117689.

Stenzel Cackowski, Emmanuel L. Barbier, Michel Dojat, and Thomas Christen. Imunity: A generalizable vae-gan solution for multicenter mr image harmonization. Medical Image Analysis, 88:102799, 2023. doi: 10.1016/j.media.2023.102799.

Jun-Yan Zhu, Taesung Park, Phillip Isola, and Alexei A. Efros. Unpaired image-to-image translation using cycle-consistent adversarial networks. In Proceedings of the IEEE International Conference on Computer Vision, pages 2223–2232, 2017.

Yunjey Choi, Youngjung Uh, Jaejun Yoo, and Jung-Woo Ha. Stargan v2: Diverse image synthesis for multiple domains. In Proceedings of the IEEE/CVF Conference on Computer Vision and Pattern Recognition, pages 8188–8197, 2020.

Siyuan Liu and Pew-Thian Yap. Learning multi-site harmonization of magnetic resonance images without traveling human phantoms. Communications Engineering, 3:6, 2024. doi: 10.1038/s44172-023-00140-w.

Farzad Beizaee, Gregory A. Lodygensky, Chris L. Adamson, Deanne K. Thompson, Jeanie L. Y. Cheong, Alicia J. Spittle, Peter J. Anderson, Christian Desrosiers, and Jose Dolz. Harmonizing flows: Leveraging normalizing flows for unsupervised and source-free mri harmonization. Medical Image Analysis, 101: 103483, 2025. doi: 10.1016/j.media.2025.103483.

Mengqi Wu, Minhui Yu, Shuaiming Jing, Pew-Thian Yap, Zhengwu Zhang, and Mingxia Liu. Unpaired volumetric harmonization of brain mri with conditional latent diffusion. Medical Image Analysis, 107: 103849, 2026. doi: 10.1016/j.media.2025.103849.

Daniel Scholz, Ayhan Can Erdur, Robbie Holland, Viktoria Ehm, Jan C. Peeken, Benedikt Wiestler, and Daniel Rueckert. Contrastive anatomy–contrast disentanglement: A domain-general mri harmonization method. In Medical Image Computing and Computer Assisted Intervention – MICCAI 2025, volume 15965, pages 100–110. Springer, 2025. doi: 10.1007/978-3-032-04978-0_10.

Fengling Hu, Andrew A. Chen, Hannah Horng, Vishnu Bashyam, Christos Davatzikos, Aaron Alexander-Bloch, Mingyao Li, Haochang Shou, Theodore D. Satterthwaite, Meichen Yu, and Russell T. Shinohara. Image harmonization: A review of statistical and deep learning methods for removing batch effects and evaluation metrics for effective harmonization. NeuroImage, 274:120125, 2023. doi: 10.1016/j.neuroimage.2023.120125.

Lianrui Zuo, Blake E. Dewey, Yihao Liu, Yufan He, Scott D. Newsome, Ellen M. Mowry, Susan M. Resnick, Jerry L. Prince, and Aaron Carass. Unsupervised mr harmonization by learning disentangled representations using information bottleneck theory. NeuroImage, 243:118569, 2021. doi: 10.1016/j.neuroimage.2021.118569.

Lianrui Zuo, Yihao Liu, Yuan Xue, Blake E. Dewey, Samuel W. Remedios, Savannah P. Hays, Murat Bilgel, Ellen M. Mowry, Scott D. Newsome, Peter A. Calabresi, Susan M. Resnick, Jerry L. Prince, and Aaron Carass. Haca3: A unified approach for multi-site mr image harmonization. Computerized Medical Imaging and Graphics, 109:102285, 2023. doi: 10.1016/j.compmedimag.2023.102285.

Blake E. Dewey, Lianrui Zuo, Aaron Carass, Yufan He, Yihao Liu, Ellen M. Mowry, Scott D. Newsome, Jiwon Oh, Peter A. Calabresi, and Jerry L. Prince. A disentangled latent space for crosssite mri harmonization. In Medical Image Computing and Computer Assisted Intervention – MICCAI 2020, volume 12267 of Lecture Notes in Computer Science, pages 720–729. Springer, 2020. doi: 10.1007/978-3-030-59728-3_70.

Jonathan Ho, Ajay Jain, and Pieter Abbeel. Denoising diffusion probabilistic models. In Advances in Neural Information Processing Systems, volume 33, pages 6840–6851, 2020.

Jiaming Song, Chenlin Meng, and Stefano Ermon. Denoising diffusion implicit models. In International Conference on Learning Representations, 2021.

Robin Rombach, Andreas Blattmann, Dominik Lorenz, Patrick Esser, and Björn Ommer. High-resolution image synthesis with latent diffusion models. In Proceedings of the IEEE/CVF Conference on Computer Vision and Pattern Recognition, pages 10684–10695, 2022. doi: 10.1109/CVPR52688.2022.01042.

Alicia Durrer, Julia Wolleb, Florentin Bieder, Tim Sinnecker, Matthias Weigel, Robin Sandkuehler, Cristina Granziera, Özgür Yaldizli, and Philippe C. Cattin. Diffusion models for contrast harmonization of magnetic resonance images. In Medical Imaging with Deep Learning, volume 227 of Proceedings of Machine Learning Research, pages 526–551. PMLR, 2024. URL https://proceedings.mlr.press/v227/durrer24a.html.

Ting Chen, Simon Kornblith, Mohammad Norouzi, and Geoffrey Hinton. A simple framework for contrastive learning of visual representations. In Proceedings of the 37th International Conference on Machine Learning, volume 119 of Proceedings of Machine Learning Research, pages 1597–1607. PMLR, 2020.

Prannay Khosla, Piotr Teterwak, Chen Wang, Aaron Sarna, Yonglong Tian, Phillip Isola, Aaron Maschinot, Ce Liu, and Dilip Krishnan. Supervised contrastive learning. In Advances in Neural Information Processing Systems, volume 33, pages 18661–18673, 2020.

Tiankai Hang, Shuyang Gu, Chen Li, Jianmin Bao, Dong Chen, Han Hu, Xin Geng, and Baining Guo. Efficient diffusion training via min-snr weighting strategy. In Proceedings of the IEEE/CVF International Conference on Computer Vision, pages 7441–7451, 2023. doi: 10.1109/ICCV51070.2023.00684.

Jacob A. Macdonald, Zhe Zhu, Brandon Konkel, Maciej A. Mazurowski, Walter F. Wiggins, and Mustafa R. Bashir. Duke liver dataset: A publicly available liver mri dataset with liver segmentation masks and series labels. Radiology: Artificial Intelligence, 5(5):e220275, 2023. doi: 10.1148/ryai.220275.

Tugba Akinci D’Antonoli, Lucas K. Berger, Ashraya K. Indrakanti, Nathan Vishwanathan, Jakob Weiss, Matthias Jung, Zeynep Berkarda, Alexander Rau, Marco Reisert, Thomas Kuestner, Alexandra Walter, Elmar M. Merkle, Daniel T. Boll, Hanns-Christian Breit, Andrew Phillip Nicoli, Martin Segeroth, Joshy Cyriac, Shan Yang, and Jakob Wasserthal. Totalsegmentator mri: Robust sequence-independent segmentation of multiple anatomic structures in mri. Radiology, 314(2):e241613, 2025. doi: 10.1148/radiol.241613.

